# Computational nanobody design using graph neural networks and Metropolis Monte Carlo sampling

**DOI:** 10.1101/2025.06.08.658414

**Authors:** Lei Wang, Xiaoming He, Gaoxing Guo, Xinzhou Qian, Qiang Huang

**Affiliations:** State Key Laboratory of Genetics and Development of Complex Phenotypes, Shanghai Engineering Research Center of Industrial Microorganisms, MOE Engineering Research Center of Gene Technology, School of Life Sciences, Fudan University, Shanghai 200438, China; Multiscale Research Institute of Complex Systems, Fudan University, Shanghai 201203, China

**Author notes:** Corresponding author at: School of Life Sciences, Fudan University, Shanghai 200438, China. These two authors contributed equally (Lei Wang, Xiaoming He).

**Keywords:** protein therapeutics, protein design, protein-protein binding, binding free energy, Metropolis algorithm, TL1A

## Abstract

Nanobodies have emerged as promising protein therapeutics due to their high-stability, low immunogenicity, and ease of production. However, experimental screening of high-affinity nanobodies for specific antigens and their post optimization remain costly and time-consuming, mainly due to the large number of possible variants. Here, we developed a computational approach that integrates graph neural networks (GNNs) with Monte Carlo Metropolis algorithm for nanobody design. We constructed a GNN model, AiPPA, to predict the protein-protein binding free energy (BFE) without requiring the complex structure, achieving a Pearson correlation of 0.62 on benchmark. We then combined AiPPA with Metropolis importance sampling to design low-BFE nanobodies from a non-affinity template. We applied this method to the antigen TL1A, and generated two affinity nanobodies. This work establishes a physics-informed deep learning method for computational nanobody design, providing a novel development strategy for protein therapeutics.

## Introduction

Antibodies have played an important role in the treatment of many diseases in recent years^1,2^. Since 2020, approximately 10 antibody-based biologics have been approved each year, bringing the total number of FDA-approved antibody therapeutics to more than 130 by the end of 2023^3^. Antibodies also dominate the pharmaceutical market: 6 of the top 10 best-selling drugs worldwide in 2023 were antibodies, with Keytruda leading the global sales^4^. The success of antibody therapeutics has driven the development of various novel types of antibodies^5^. Among these, nanobody (Nb), as the smallest antigen-binding fragment^6^ (∼15 kDa, one-tenth the size of traditional antibodies), offers remarkable tissue and tumour penetration^7,8^. The small size, high stability^9^, low immunogenicity^10^, and ease of production^11^ has further solidified its role in therapeutic innovation. To date, three Nb-based therapeutics - Caplacizumab^12^, Envafolimab^13^, and Ozoralizumab^14^ - have been approved for clinical use; and, many more are in clinical trials^15^. Thus, nanobodies hold great potential for the treatment of a wide range of diseases.

Nanobodies are primarily discovered through library screening; however, these screened nanobodies often exhibit low binding affinity to disease-related antigens^16,17^. High affinity is essential not only to enhance the efficacy of therapeutic nanobodies but also to reduce off-target effects and minimize potential side effects^18^. Therefore, affinity maturation or optimization for the screened nanobodies is often required. Two primary approaches have been developed to improve the binding affinities of nanobodies *in vitro*. The first is based on the hypermutagenic properties of specific cells, such as B cells or H1229 cells, and to replicate the *in vivo* hypermutation process in an *in vitro* setting^19,20^. However, this method heavily relies on cells types and requires complicated experimental manipulations^18^. The second approach involves generation of mutant libraries by introducing random mutations into nanobodies, followed by selection systems to screen for variants with increased affinities^21^. Due to the randomness of the mutations, this method often requires large libraries and multiple rounds of screening, making it costly and time-consuming^22,23^. In addition, *in vitro* affinity maturation often has the risk to compromise the stability and other properties of the nanobodies^24,25^.

To address the challenges of *in vitro* affinity maturation, computational design methods have been developed to improve efficiency and reduce costs^21,22^. These methods typically treat affinity optimization of a low-affinity antibody or nanobody as an energy minimization task, aiming to design or generate sequence variants with lower binding free energies (i.e., higher binding affinities). This process usually involves two steps: low-energy sequence search in a large sequence space and binding affinity evaluation of the searched sequences, with the latter being more critical for a successful computational optimization. However, existing computational affinity evaluation methods, such as PRODIGY^26^, often rely on the accurate antibody-antigen complex structures. Even with advanced prediction tools like AlphaFold-Multimer^27^ or AlphaFold3 (AF3)^28^, predicting protein-protein complex structures remain time-consuming for large numbers of antibody variants, and sometimes the predictions are also insufficiently accurate for the binding free energy calculations^29,30^. Consequently, protein-protein complex structure-based methods are difficult to deal with large numbers of potential affinity variants, often resulting in limited affinity improvement^31,32^. Thus, there is still a need for more efficient computational methods to identify affinity variants of antibodies and nanobodies targeting given antigens.

In this study, we considered nanobody design for a specific antigen as an *in silico* affinity maturation or optimization process, staring from a non-affinity template nanobody. Based on this, we introduced a computational design framework that integrates graph neural networks (GNNs) with Markov chain Monte Carlo (MC) Metropolis sampling. First, we constructed a GNN model called AiPPA (Ai-enabled Protein-Protein Affinity predictor) for accurate prediction of the thermodynamic protein-protein binding free energy (BFE) without the need for their complex structures. This model was validated by the Kastritis benchmark. Then, we used the MC Metropolis importance sampling algorithm^33^ to stochastically explore sequence variants within the complementarity-determining regions (CDRs) of the template. During sampling, AiPPA calculated the binding free energies of the nanobody variants to the antigen, guiding the exploration of low-BFE sequences based on the Boltzmann factor of free energy changes. We applied this physics-informed deep learning approach to design nanobodies targeting TL1A (Tumor necrosis factor-like cytokine 1A), an antigen implicated in autoimmune diseases. Experimental validation using bio-layer interferometry (BLI) confirmed that two of the six designed nanobodies bind TL1A with equilibrium dissociation constants of ∼2.0 µM, demonstrating the effectiveness of our design method.

## Results

### 1. Architecture and training of AiPPA

In protein-protein binding affinity prediction, various machine learning-based models have been developed^26,34,35^. Among them, structure-based models usually use the complex structures to construct structural features from the protein-protein interaction (PPI) interfaces and non-interfaces, and then apply linear regression or support vector regression to fit the protein-protein binding affinities (or binding free energies). To avoid using the protein-protein complex structure, in this study, we used GNNs to develop AiPPA that models two interacting protein partners (A and B) as two separate graphs (Fig. 1A). Then, the graph representation learning was applied to each protein graph (A or B); then, these two protein graphs are combined to predict the BFE of proteins A and B (Fig. 1B). In this way, AiPPA enables a fast prediction of the binding free energy without the need for the accurate A-B complex structure, providing a solid basis for the subsequent large-scale design of the low-BFE variants from a given template nanobody.

**Figure 1.**
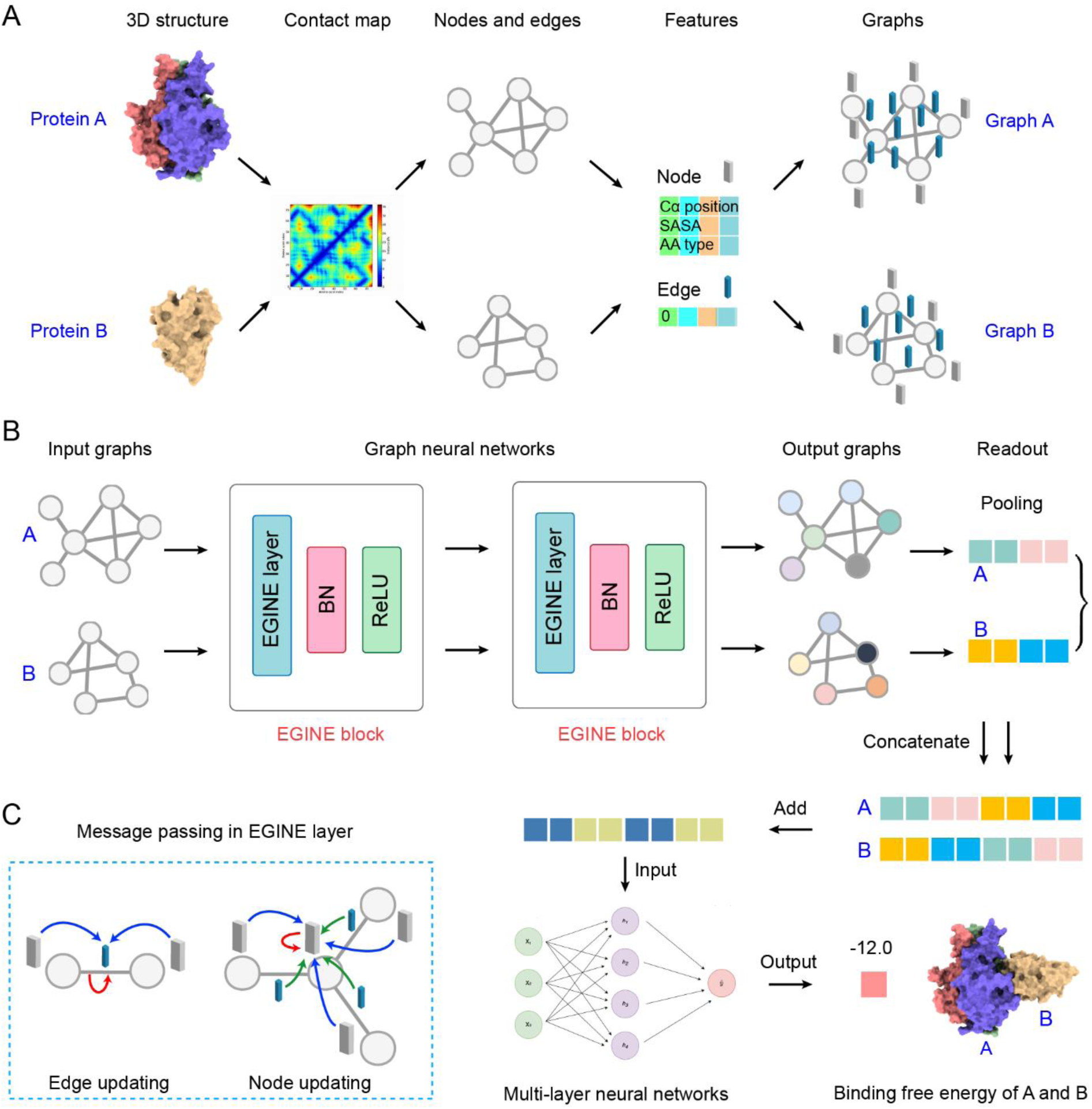
Architecture of the AiPPA model. (A) Graph representation of two interacting proteins (A and B). Residues of the proteins serve as nodes, with their physicochemical properties encoded as node features. Edges are defined by the contact map and represent spatially adjacent amino acid residues based on the 3D structures of the proteins. The initial edge features are set to 0. (B) Architecture and learning process of the AiPPA model. The AiPPA model consists of two serially connected EGINE blocks, which iteratively update the graph representations of proteins based on the training data. The graph representations of the two proteins are then combined using concatenation and addition operations and fed into a MLP to calculate the binding free energy between A and B. (C) Message passing in the EGINE blocks. During the learning process, message is passing between the nodes and edges of the proteins, and iterative updates. Blue arrows indicate messages from adjacent edges, green arrows represent messages from neighboring nodes, and red arrows denote self-referential messages from edges or nodes.

As shown in Fig. 1A, residues were represented as nodes in the protein graph (A or B), and the initial values of their features were assigned according to three properties that are closely related to the binding free energy: C_α_ coordinates, solvent-accessible surface area (SASA), and residue type. Among them, the C_α_ coordinates characterize the positions, SASA reflects the exposure of a residue to the solvent, since a surface residue is more likely to form the protein-protein contacts; and the residue type indicates the side chains that contribute significantly to the residue interactions. To facilitate message passing between the spatially close residues, we also added edges to the protein graphs for any two nodes whose C_α_-C_α_ distance is less than a given cutoff value, *θ*. After several tests, we found that AiPPA achieved optimal performance at *θ* = 8 Å (Fig. S1A). Since the edges represent the relationships between residues and support the message passing between the nodes, we ever defined the initial edge features according to the distances between two nodes, etc. During the model training process, however, the evaluation of the impact of various feature combinations on the model performance showed that the node features contributed significantly to the model performance, but the edge features did not (Fig. S1B). Thus, the initial edge features in AiPPA were set as zero vectors.

As shown in Fig. 1B, the core of AiPPA consists of two EGINE blocks, which are based on the Graph Isomorphism Networks (GINs), a powerful GNN that uses an injective aggregation function and has the graph classification ability approaching the Weisfeiler-Lehman test^36^. Hu et al. introduced edge features into the aggregation process and thereby developed GINE (GIN with edges), which further improved its performance^37^. In this study, we further introduced “edge updating” to GINE to construct the edge-updating GINE (EGINE), in order to improve the message passing between the nodes and the edges (Fig. 1C). Then, an EGINE layer combines a batch normalization (BN) layer and a ReLU activation function layer to form an EGINE block (Fig. 1B). After the processing by two stacking EGINE blocks, the output edge features of the two graphs formed the final representations of the proteins A and B. These two representations were then merged via concatenate and add operations, and then fed into a multilayer perceptron (MLP), which finally outputs the binding free energy of the proteins A and B (Fig. 1B).

To train AiPPA, we screened a dataset of 1,858 PPI samples from the PDBbind database (v2020)^38^(see Supplementary Methods and Fig. S2), which was the largest training dataset available for the protein-protein binding affinity modeling at the time of this study. We divided this dataset into training and validation sets in an 8:2 ratio for the model training and validation, respectively, with the similar energy distributions (Fig. S3A). As the binding free energy prediction is a regression task, we evaluated the model performance using the Pearson correlation coefficient (PCC) and root mean square error (RMSE). Training of the AiPPA model was stopped when the validation loss reached its minimum, in order to avoid the model overfitting (Fig. S3B).

### 2. Model performance of AiPPA

To verify the AiPPA model, we used a popular benchmark for the protein-protein binding affinities: the Kastritis benchmark^39^, which was introduced by Kastritis et al. in 2011 and contains 144 protein-protein complexes with experimentally determined binding affinities and high-resolution structures. This dataset spans a wide range of biological functions such as antibody-antigen and enzyme-inhibitor interactions, and covers a wide range of binding affinities, with values of equilibrium dissociation constant (*K*_d_) ranging from 10^-14^ to 10^-5^ M. Thus, the Kastritis benchmark has been widely used to evaluate various affinity prediction models. For a reliable comparison with other representative models, we selected 138 samples from this benchmark as the test set (see Supplementary Methods). The free energy distributions of this test set are similar as those of the training and validation sets, allowing a reliable comparison of the model performance (Fig. S3A).

During the period of our study, three main binding affinity prediction models were available for the performance comparison: PPA_Pred2^35^, PPI-Affinity^34^, and PRODIGY^26^. Among them, PPA_Pred2 uses protein sequences as input and was trained by 453 dimers selected from the PDBbind (v2015) database and categorized into six functional classes. PPI-Affinity was trained by 833 samples selected from PDBbind (v2020), and used 26 structure-based molecular descriptors and support vector machine (SVM) to predict the binding affinities. Both models offer web interfaces for the prediction. Unlike them, PRODIGY was trained by the Kastritis benchmark, and used six structural features from the PPI and non-interaction interfaces to predict the affinities. Also, this model is available as a standalone program, and thus is the most widely used model.

Based on the test set, we calculated the PCC and RMSE values for each model. As shown in Fig. 2A and Table S1, AiPPA achieved the best PCC of 0.62, which was 11%, 26%, and 15% higher than those of PPA_Pred2, PRODIGY, and PPA-Affinity, respectively; AiPPA also had a relatively low RMSE (Fig. 2A). Thus, the predicted affinities by AiPPA showed a strong correlation with the experimental values (Fig. 2B). Although PPI-Affinity had the lowest RMSE value, its correlation was not good (PCC = 0.54). This suggests that AiPPA has the best ability to rank the binding free energies of the samples, an important property when searching for the lowest energy sequences from large numbers of possible sequences. Significantly, AiPPA performed well for those samples with binding free energies around -10 kcal·mol^-1^, the most frequent region in the energy distributions of the training set (Fig. S3A).

**Figure 2.**
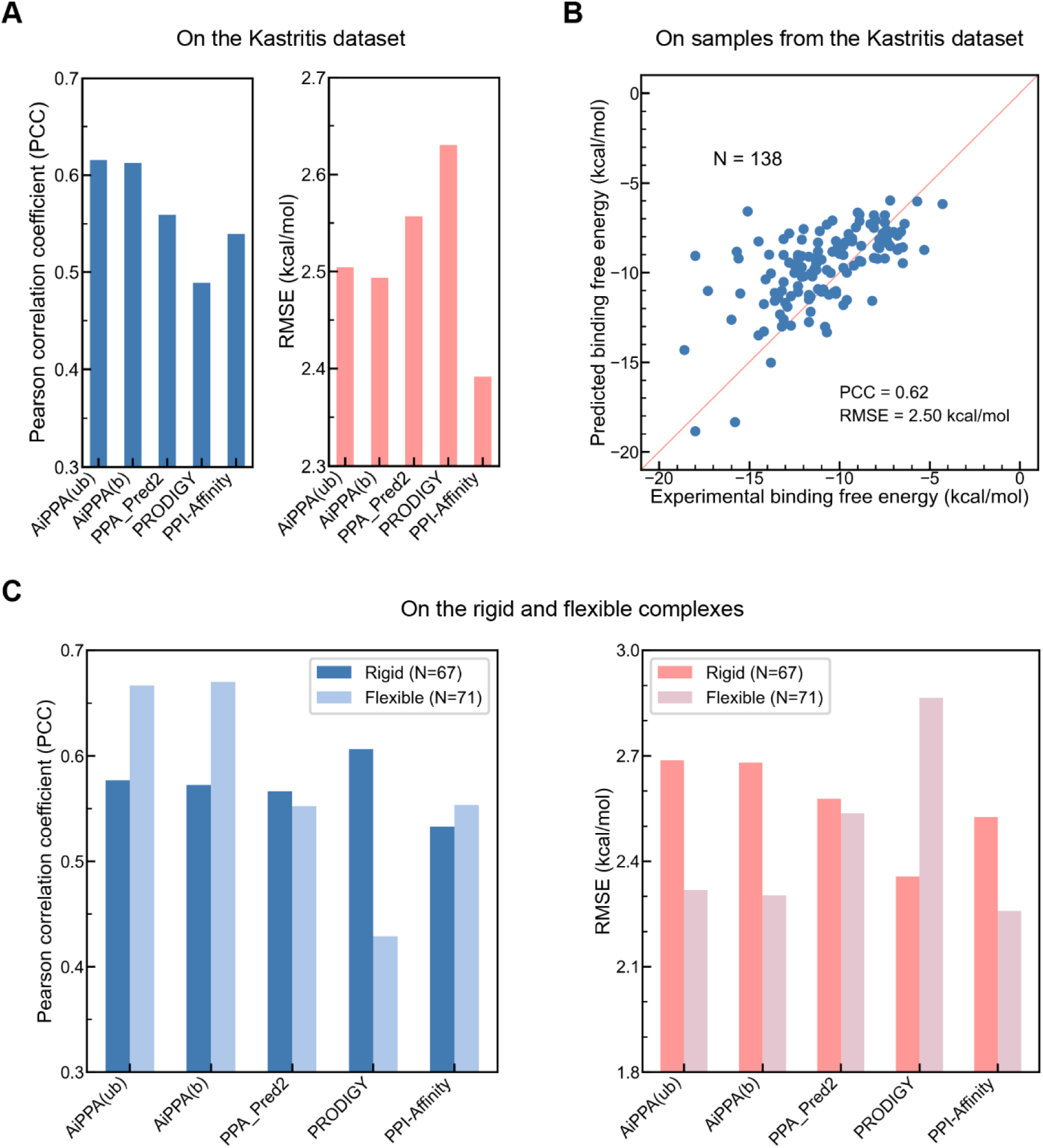
Performance of the AiPPA model. (A) Comparison of AiPPA, PPA_Pred2, PRODIGY, and PPI-Affinity on the Kastritis benchmark (test set). The left panel shows the Pearson correlation coefficient (PCC), with higher values indicating better predictive accuracy, while the right panel displays the root mean square error (RMSE), where lower values correspond to smaller deviations between predicted and experimental values. (B) Correlation plot of predicted versus experimental binding free energies of the test set. (C) Performance metrics (PCC, left; RMSE, right) for AiPPA, PPA_Pred2, PRODIGY, and PPI-Affinity on rigid and flexible subsets of the test set.

As known, proteins usually undergo conformational changes during their binding processes. Therefore, to accurately predict the interactions between two “flexible” proteins are more difficult than those between two relatively rigid proteins. To evaluate the effectiveness of AiPPA for this kind of interactions, we further divided the test set into rigid complexes (Rigid, I_RMSD ≤ 1.0 Å, *N* = 67, Fig. S4A) and flexible complexes (Flexible, I_RMSD > 1.0 Å, *N* = 71, Fig. S4B) based on Vangone and Bonvin’s criteria^26^, and then assessed the model performance on these two subsets (Fig. 2C and Table S1). As seen, sequence-based model PPA_Pred2 exhibited similar performance for both subsets, because the input sequence features were the same for both cases. The highest PCC and the lowest RMSE indicated that PRODIGY had the best prediction for the rigid complexes. However, for the flexible complexes, its performance dropped by ∼30% in the PCC, implying that the used molecular descriptors were not enough for the accurate prediction of the binding free energies in the flexible interactions. In contrast, PPI-Affinity and AiPPA performed better on the flexible complexes; significantly, AiPPA achieved the best results on both subsets, with a high accuracy for the flexible complexes (PCC = 0.67), outperforming other three models.

For the flexible complexes, conformational changes at the binding interface often result in differences between the bound and unbound states of the protein (A or B; Fig. S4B). To evaluate the robustness of AiPPA to such conformational variability, we assessed its predictions using both bound and unbound protein conformations, respectively. As shown in Fig. 2C, AiPPA demonstrated consistent performance compared to Fig. 2A, indicating its ability to accurately predict binding free energies despite conformational changes during binding. This capability is critical for its application in computational nanobody design, where flexibility at the interface is a key consideration.

### 3. Monte Carlo sampling of CDR variants

A typical nanobody comprises four framework fragments (FR1, FR2, FR3 and FR4) and three CDR loops (Fig. 3A). The nanobody frameworks are very conserved, but the CDR loops are not; indeed, the CDRs play a crucial role in the binding to the antigens^40^. As a result, nanobody design and engineering usually focus on the design of the CDR loop sequences, especially that of the CDR3 loop. In order to integrate AiPPA, as mentioned, we considered the nanobody design for a specific antigen as an *in silico* generation of the CDR variants of a non-affinity template nanobody, especially a nanobody with a humanized framework. To this end, we employed MC simulation, a thermodynamically rigorous method rooted in statistical physics and widely used for sampling complex energy landscapes, to explore CDR variants of the template. Using AiPPA, we calculated the binding free energy of each sampled variant with the target and accepted or rejected variants based on the Metropolis criterion, which ensures convergence to a Boltzmann-weighted equilibrium distribution^41^. As illustrated in Fig. 3B, each MC iteration comprises five steps: (1) generating a new CDR variant from the previous sequence; (2) predicting its solubility index; (3) predicting its 3D structure; (4) evaluating its binding free energy to the target using AiPPA; and (5) accepting or rejecting the variant according to the Boltzmann-weighted energy change criterion. This physics-based framework enables efficient exploration of high-dimensional sequence design spaces, guiding the search toward low-BFE, high-affinity nanobody variants.

**Figure 3.**
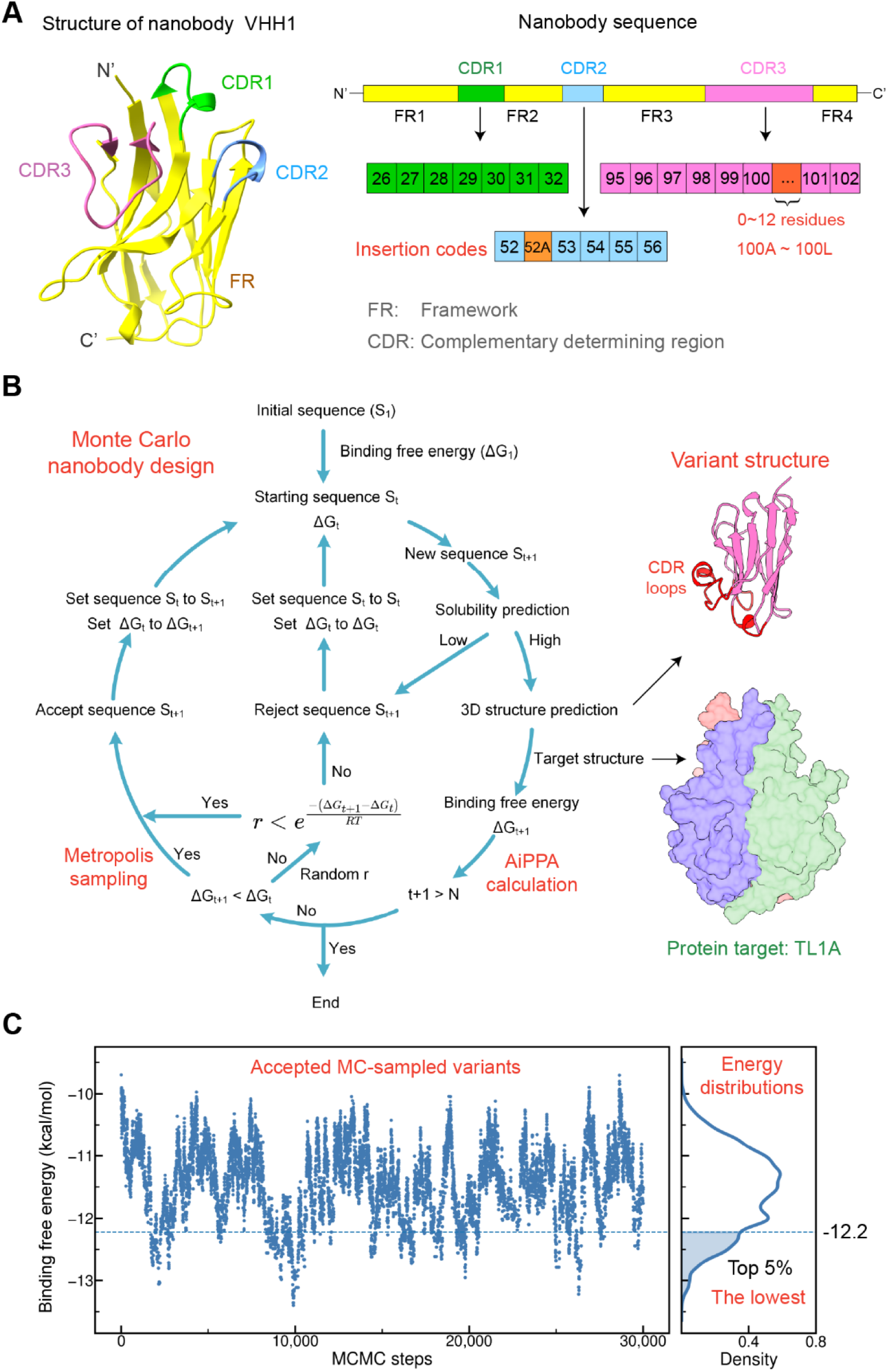
Design of high-affinity nanobodies by combining AiPPA and MC Metropolis sampling. **(A)** 3D structure and sequence of the template nanobody VHH1. The framework is shown in yellow, with highlighted CDRs: CDR1 (green), CDR2 (blue), and CDR3 (pink). During the design process, CDR2 and CDR3 lengths were varied by inserting amino acids at specific positions: position 52A for CDR2 and positions 100A–100L for CDR3, depending on the sequence length. (B) MC sampling for low-BFE nanobody design based on the Metropolis Boltzmann-weighted criterion. Variants are iteratively generated through MC sampling. Each iteration involves: (1) generating a new sequence (S_t+1_) via random mutation, followed by solubility prediction; low-solubility sequences are discarded and regenerated; (2) predicting the 3D structure of high-solubility sequences and calculating their binding free energy (ΔG_t+1_) with the target protein using AiPPA; and (3) applying the thermodynamic Metropolis criterion to compare ΔG_t+1_ with the previous iteration’s binding free energy (ΔG_t_) to determine sequence acceptance or rejection. The variant sequence and binding free energy are updated iteratively. (C) Accepted variants and their binding free energies from 30,000 MC sampling iterations. The top 5% of variants exhibited binding free energies below -12.2 kcal·mol⁻¹, as indicated by the blue dashed line.

To enhance the sampling efficiency for nature-like variants, we analyzed ∼20,000 natural nanobody sequences from the INDI database^42^ to extract sequence features in their CDRs, defined by the Chothia numbering (Fig. 3A, right; Table S2). After removing duplicates, we obtained 3,108 CDR1, 3,812 CDR2, and 10,770 CDR3 loop sequences, which were used to analyze length and amino acid distributions (Fig. S5). Our analysis revealed that the most frequent lengths for CDR1, CDR2, and CDR3 are 7, 5–6, and 5–20 residues (> 90% frequency), respectively. Additionally, specific amino acid preferences were observed at certain positions, such as glycine (G) at H26 and tyrosine (Y) at H102, suggesting conservation in CDR loop sequences. To generate nature-like variants during MC sampling, we constrained CDR lengths to these ranges: CDR1 = 7 aa, CDR2 = 5–6 aa, and CDR3 = 5–20 aa. Length transition probabilities were calculated based on current CDR lengths (Methods; Fig. S6). Furthermore, amino acid sampling probabilities were weighted according to their natural distribution frequencies in the CDR loops.

Note that, to ensure a soluble expression of a sampled variant, in each MC iteration step we used Protein-Sol^43^ to predict the solubility index of the variant and rejected those with a solubility index < 0.45. For a variant with a solubility index ≥ 0.45, its 3D structure was then predicted using IgFold^44^, which provides similar accuracy to AlphaFold2^45^ but is almost 60 times faster. The binding free energy of the variant to the protein target was then calculated using AiPPA to determine its probability of acceptance using the Metropolis importance sampling algorithm (Methods, Fig. 3B). The MC affinity variant sampling process was terminated when the maximum MC iteration steps of 30,000 were reached.

### 4. Low-BFE variant design from a template nanobody

To validate our method, we applied it to design low-BFE nanobodies targeting TL1A, a key therapeutic target for inflammatory bowel diseases (IBDs) such as ulcerative colitis and Crohn’s disease, which remain among the most challenging diseases of the 21st century^46,47^. In this study, we used the nanobody VHH1 as the template for the design (Fig. S7). VHH1 is an affinity nanobody targeting TNF-α, which is a homologue protein of TL1A with relevance to rheumatoid arthritis (RA) and IBDs^48^ (sequence in Fig. S7A). We expressed and then confirmed via bio-layer interferometry (BLI) that VHH1 does not bind TL1A (Fig. S7B, C). Starting from the VHH1 sequence, we performed 30,000 MC simulation steps, sampling approximately 9,400 variants accepted by the Metropolis algorithm (Fig. 3C). Notably, ∼3,200 variants exhibited predicted binding free energies below -12.0 kcal·mol⁻¹, with 34 variants reaching energies < - 13.0 kcal·mol⁻¹ and a minimum of -13.4 kcal·mol⁻¹. These results demonstrate the ability of our MC sampling framework (Fig. 3b) to design nanobodies with potential TL1A-binding affinity. For further evaluation, we selected the top 5% of variants ranked by binding free energy (ΔG < -12.2 kcal·mol⁻¹; Fig. 3C).

To evaluate the sequence uniqueness and diversity of the top 5% of low-energy variants, we calculated the Levenshtein distances (LDs) between their CDRs and those of VHH1, as well as the shortest LDs between the generated variants and natural nanobodies. As shown in Fig. 4a, most variants exhibited LDs > 3 for CDR1 and CDR2 relative to VHH1, while CDR3 showed a minimum LD of 10, with most distances ranging between 13 and 15 (Fig. 4A, right), indicating significant divergence from VHH1. Furthermore, the shortest LDs indicated that CDR1 and CDR2 exhibited a high sequence similarity to those of natural nanobodies (LD = 1-2), with some variants matching natural sequences exactly (LD = 0). In contrast, CDR3 loops displayed a minimum LD of 6 and a maximum of 11 compared to the natural CDR3 loops (Fig. 4A, left), highlighting both the critical role of CDR3 in binding affinity and the potential for generating novel nanobody sequences. To further assess diversity, we computed pairwise LDs between the CDR loops of the variants. As shown in Fig. 4B, the majorities of LDs were non-zero, demonstrating the high sequence diversity of the low-energy variants sampled by the MC simulation.

**Figure 4.**
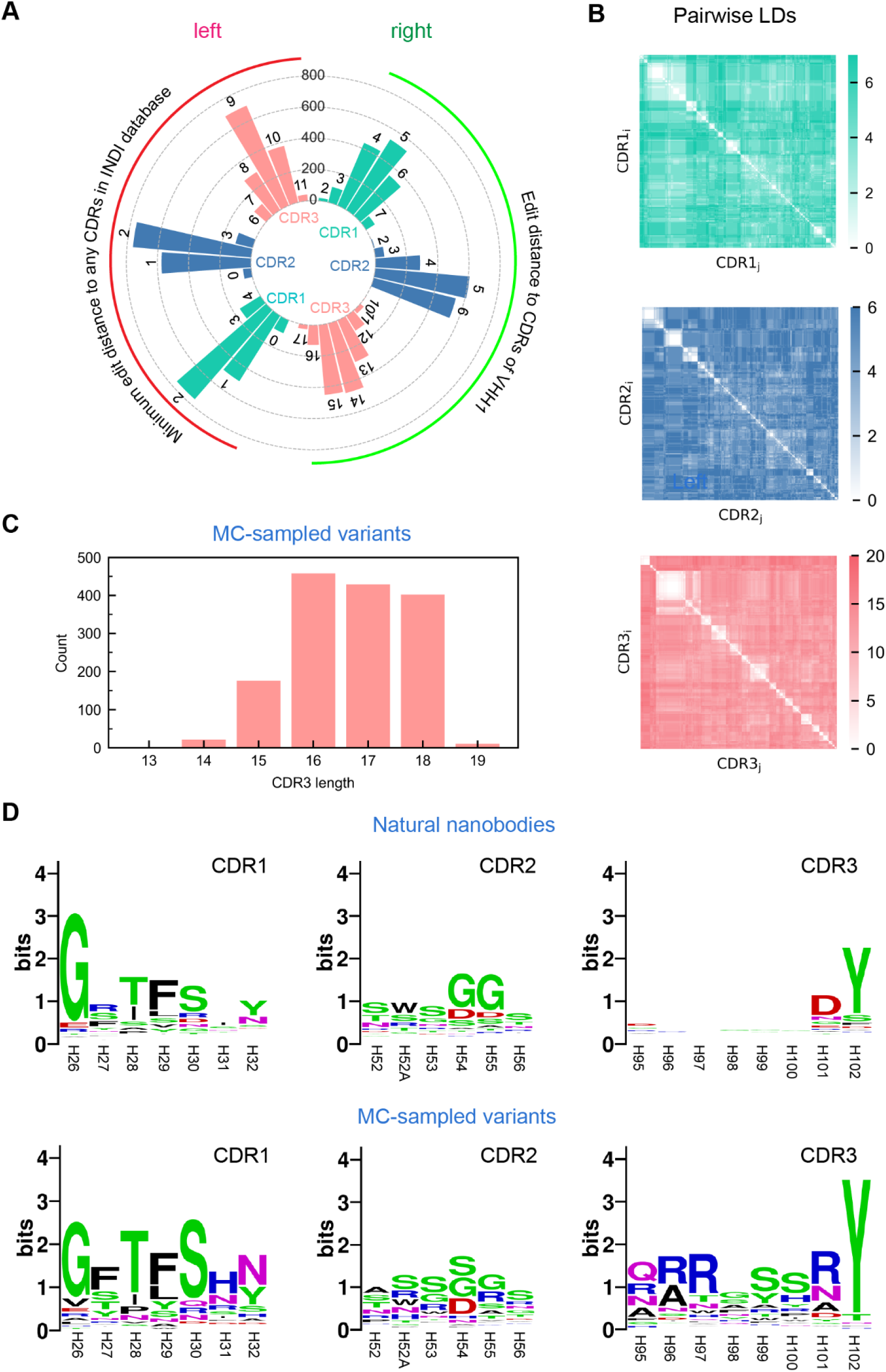
Sequence characteristics of the MC-sampled affinity nanobodies targeting TL1A. (A) Levenshtein distances (LDs) between the CDRs of the nanobodies and the template VHH1 (right) and the minimum LDs between the CDRs of the nanobodies and natural nanobodies (left). Larger edit distances indicate greater sequence divergence. (B) Pairwise LDs among the CDR loops of the sampled nanobodies. (C) Length distribution of the CDR3 loops in the sampled nanobodies. (D) Amino acid distribution logo plots for key positions in the CDR loops of natural nanobodies and the sampled nanobodies. Logo plots were generated using WebLogo3^58^.

To investigate whether the sampled variants exhibited specific sequence features for TL1A binding, we analyzed the length distribution of CDR3 loops and the amino acid composition of all three CDR loops. As shown in Fig. 4C, the CDR3 loops of the variants were predominantly longer (16-18 amino acids) compared to the 14-residue CDR3 of VHH1, suggesting that increased loop length may expand the binding interface with TL1A, a potential determinant of high affinity. Sequence logos revealed that although the dominant amino acids in CDR1 and CDR2 were comparable to those of VHH1, their relative frequencies differed significantly (Fig. 4D). Notably, CDR3 loops displayed a strong preference for specific residues, particularly arginine (R) and serine (S), which may enhance TL1A binding through electrostatic interactions and hydrogen bonding. These findings demonstrate that the MC sampling method has captured the sequence features critical for designing low-BFE nanobodies targeting TL1A.

### 5. Site-specific filtering of designed nanobodies

Because Nb-TL1A complex structures were not used in the process of the MC variant generation, we did not know the exact binding site of a generated variant on TL1A. Although the top 5% variants in above were predicted as potential affinity variants, their actual binding sites on TL1A might be different. To identify those variants that bind to a desired binding site of TL1A (i.e., residues related to DcR3 binding^49^) (Fig. 5A, top), we performed computational site-specific filtering on these variants. In addition, given the high false-positive rates in protein design^50^, a systematic procedure shown in Fig. 5A (middle) was used to filter and refine the top 5% MC-sampled variants, and finally to get the designs binding to the desired site (Fig. 5A, bottom).

**Figure 5.**
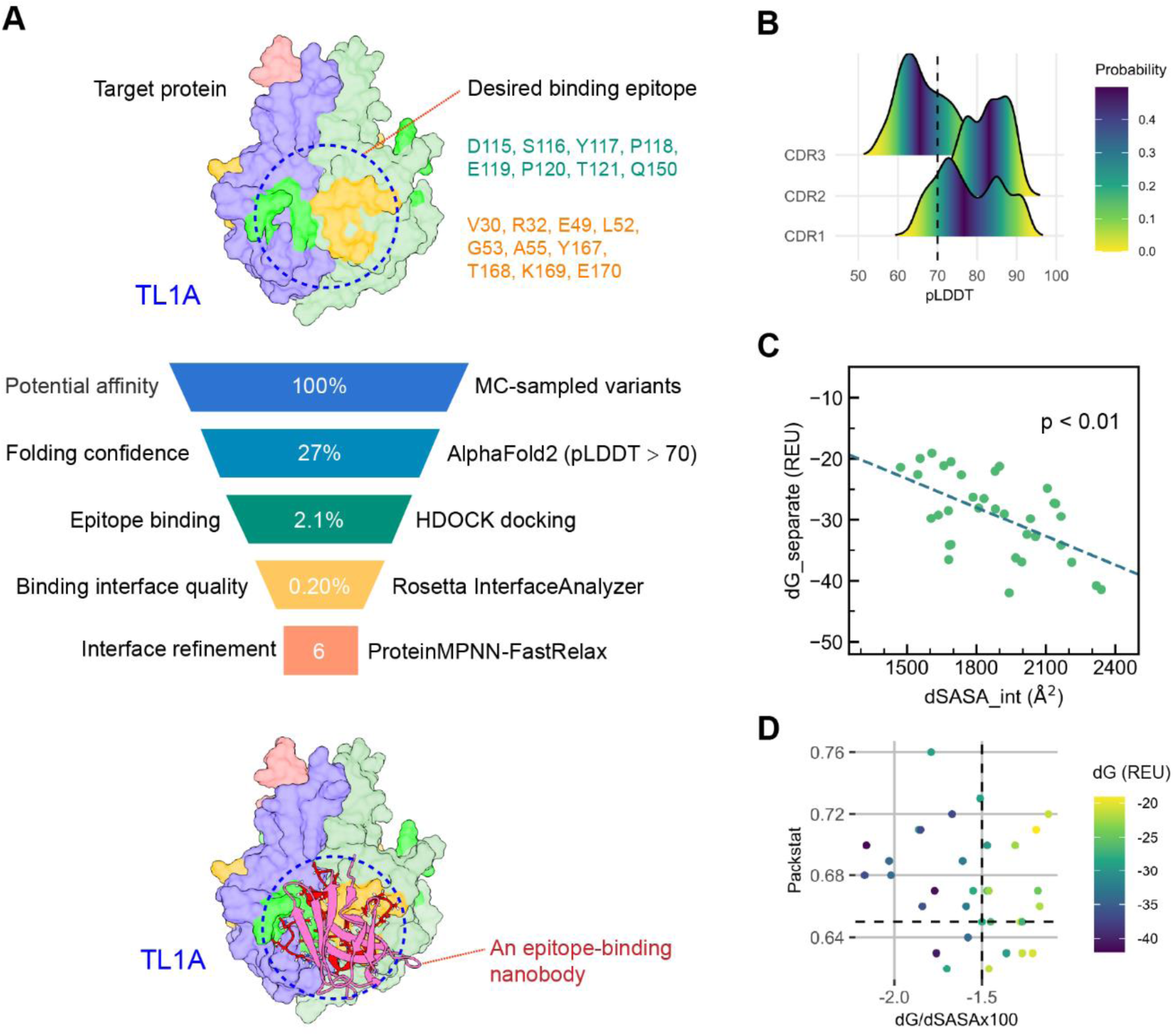
Site-specific filtering of designed nanobodies targeting TL1A. **(A)** Workflow for screening nanobodies binding to a desired binding site. The homo-trimeric TL1A is depicted with its three chains colored in purple, light green, and light pink (top and bottom panels). The target epitope, spanning two chains, is highlighted by the blue circle and the lime and orange residues, respectively (top and bottom panels). From 1,500 predicted low-binding free energy variants, 6 candidate nanobodies (0.4%) binding to the specified TL1A epitope were identified. The nanobody is shown in pink, with its CDRs displayed as red sticks (bottom panel). (B) Distributions of pLDDT scores for the CDR loops of the sampled variants. The black dashed line indicates pLDDT > 70, representing high-confidence predicted structures. (C) Relationship between binding free energy (ΔG_separated) and interface area (ΔSASA_int) in interface quality analysis. (D) Interface quality evaluation using three parameters: packstat > 0.65, ΔG/ΔSASA × 100 < -1.5, and ΔG_separated < -40. Binding free energy (in REU) is represented by a color gradient, with darker shades indicating lower (more favorable) binding energies.

We first predicted the 3D structures of all the variants using AlphaFold2 and then selected those variants with a pLDDT (predicted Local Distance Difference Test) value > 70. As shown in Fig. 5B, the pLDDT distributions of the variants indicated that the prediction confidence of the CDR1 and CDR2 loops was high, but that of the CDR3 loops was relatively low, probably due to the greater length and sequence variability. Using the threshold of pLDDT > 70 for all the CDR loops to filter the top 5% variants, and 405 matching variants were then obtained. To identify their potential binding sites on TL1A, these matching variants were docked to TL1A using the HDOCK program^51^, and the top-ranked docked pose of a given variant was used as its representative binding conformation to TL1A. This yielded 31 variants bound to the desired binding epitope, as illustrated in Fig. S8. We then minimized the folding energies of their complex structures with the Rosetta Relax protocol and then analyzed their Nb-TL1A interfaces using the InterfaceAnalyzer module.

Three interface properties from InterfaceAnalyzer were considered for the further filtering: Rosetta binding free energy (dG_separated) and solvent accessible area buried at the interface (dSASA_int), and packing state (packstat). Indeed, we found that the binding free energies and the solvent accessible areas buried at the interfaces of the 31 variants showed a linear correlation: a larger dSASA_int corresponds to a lower binding free energy (Fig. 5C). Therefore, the interface property criteria (dG_separated < -40, dG/dSASA × 100 < -1.5, and packstat > 0.65) were used to filter the remaining 31 variants, and three variants were found to meet these criteria (i.e., 0.20% of the top 5% variants).

Based on the docked complex structures of these 3 variants, we used ProteinMPNN^52^ to locally refine their interface residues in the CDR3 loops, with a preference for amino acids “YGSRV” according to the study by Yi et al.^53^. Finally, to reduce the possible number of the refined variants for experimental validation, we performed a self-consistency prediction of the Nb-TL1A complex structures for all the refined variants using AlphaFold2, and then selected those variants that bind to the desired epitope and match the interface criteria above. In this way, six refined variants were finally selected for the subsequent experimental validation, as listed in Table S3.

### 6. Experimental validation and structural modeling of designed nanobodies

To verify the binding activities of the 6 designed nanobodies, we expressed them in *E. coli* Rosetta DE3 cells. Except for the nanobody Tr4s18, the other 5 designed nanobodies were successfully expressed in the soluble form. We purified these nanobodies with the 6-His tags using nickel-based affinity chromatography, and then cleaved the 6-His tags with TEV protease. Then, we further purified the cleaved samples by reverse affinity chromatography to obtain the tag-free proteins of the designed nanobodies (Fig. S9).

To verify the binding activities of the designed nanobodies, we used BLI to measure their binding kinetics to TL1A. To this end, biotin-labelled TL1A proteins were immobilized on the SA sensors, and the nanobodies at different concentrations were then introduced as the mobile phase to observe binding and dissociation over a specific concentration range. The measurements showed that two designed nanobodies, Br3s95 and Tr3s90, exhibited clear binding curves (Fig. 6A). And the best-fitting *K*_d_ (equilibrium dissociation constant) values of the Br3s95 and Tr3s90 binding to TL1A were 2.56 and 1.86 μM, respectively. This indicates that we have successfully designed two binding nanobodies for TL1A using our physics-informed deep learning design approach.

**Figure 6.**
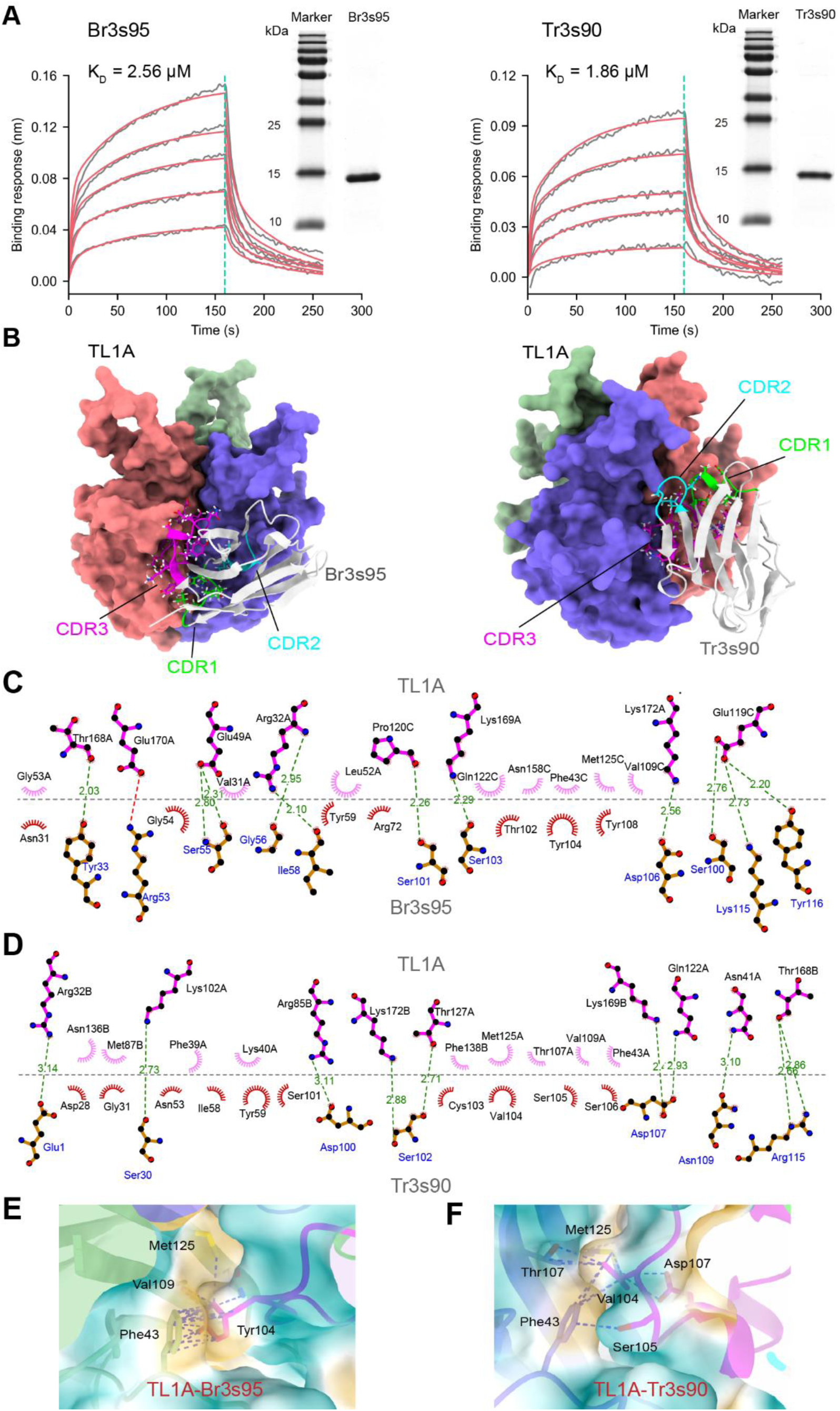
Experimental validation and structural modeling of the designed nanobodies. (A) Binding kinetics of nanobodies Br3s95 and Tr3s90 to TL1A measured by Bio-Layer Interferometry (BLI), with corresponding SDS-PAGE analysis on the right. Grey traces represent raw BLI data, and red curves show the best-fit models. (B) Atomic models of Br3s95 and Tr3s90 in complex with TL1A. TL1A is shown as a surface model, while Br3s95 and Tr3s90 are depicted as cartoon models in light grey. CDR1, CDR2, and CDR3 loops are highlighted in lime, cyan, and magenta, respectively. (C) Hydrogen bonds between the CDR loops of Br3s95 and TL1A, indicated by green dashed lines. (D) Hydrogen bonds between the CDR loops of Tr3s90 and TL1A, indicated by green dashed lines. (E) Hydrophobic interactions of Br3s95 mediated by residue F43 of TL1A. Dashed lines indicate surrounding residues in the hydrophobic core, and the nearby surface is colored by hydrophobicity (orange: hydrophobic; cyan: hydrophilic). (F) Hydrophobic interactions of Tr3s90 mediated by residue F43 of TL1A. Dashed lines indicate surrounding residues in the hydrophobic core, and the nearby surface is colored by hydrophobicity (orange: hydrophobic; cyan: hydrophilic). Panels C and D were generated using Ligplot+^59^.

In addition, since the affinity maturation process can sometimes affect the structural stability of the antibodies^24^, we further evaluated the protein stability of the designed nanobodies compared to the starting template VHH1 by determining their melting temperature (*T*_m_) using nanoscale differential scanning fluorimetry (nanoDSF). As shown in Fig. S10, the *T*_m_ value of the template VHH1 is about 70.0°C, but the *T*_m_ values of Br3s95 and Tr3s90 are about 72.3°C and 77.1°C, respectively, slightly higher than that of VHH1. This indicates that the computational design not only increased the binding affinity but also slightly improved the protein stability of the nanobodies.

To elucidate the key interactions contributing to the binding affinities of the designed nanobodies, we performed structural analysis based on the complex structures predicted by AF3. As shown in Fig. 6B, Br3s95 and Tr3s90 interact with TL1A primarily through their CDR3 loops; however, their binding poses at the interface differ: Br3s95 adopts an upward-facing conformation, whereas Tr3s90 adopts a downward-facing conformation. In Br3s95, the CDR2 and CDR3 loops form hydrogen bonds with TL1A (Fig. 6C), whereas in Tr3s90 the hydrogen bonds are predominantly formed by the CDR3 loop (Fig. 6D). For both nanobodies, serine and basic amino acids are central to the hydrogen bonding networks (Fig. 6C, D). In addition, both nanobodies interact with the hydrophobic residue F43 on the TL1A surface, which forms the core of the hydrophobic interactions. In Br3s95, F43 interacts with Y104 in the CDR3 loop through π-π stacking (Fig. 6E), whereas in Tr3s90, F43 interacts with V104 of the CDR3 loop with its surrounding residues (Fig. 6F). Apparently, besides the hydrogen bonding, these hydrophobic interactions further stabilize the Nb-TL1A binding. Therefore, hydrogen bonding and hydrophobic interactions are the main driving forces for the binding of the designed nanobodies to TL1A.

## Discussion

In this study, we have combined GNNs with Metropolis MC sampling to develop a physics-informed deep learning approach for nanobody design starting with a template nanobody. We first constructed the GNN-based model AiPPA that predicts protein-protein binding free energies without requiring structural information of the PPI complexes, achieving a Pearson correlation of 0.62 on the Kastritis benchmark. Then, we used the Metropolis importance algorithm to generate potential affinity variants of the template nanobody by calculating their binding free energies to the given target antigen with AiPPA. We used this computational method to design low-BFE nanobodies targeting the autoimmune disease-related protein, TL1A. And the BLI experiments validated that two designed nanobodies could bind TL1A, validating the effectiveness of our method for computational nanobody design.

To the best of our knowledge, AiPPA is the first structure-based GNN model that uses only the structures of two individual protein partners (A and B) to predict the protein-protein binding free energy, without the need for their complex structure. In contrast, many previous models require the protein-protein interface features extracted from the complex structures^26,34^. In practice, it is usually difficult to accurately predict the complex structures using molecular docking or deep learning models such as AF3^28,30^. Meanwhile, for large-scale computational tasks such as those in this study, accurate prediction of the complex structures by molecular docking or AF3 is often not feasible because the prediction even for a single protein-protein complex structure takes a relatively long time, up to several minutes or tens of minutes on a workstation computer with ordinary GPUs. Therefore, affinity prediction methods that do not require any protein-protein complex structures are very helpful for the large-scale evaluation of the protein-protein binding affinities, as in our study.

As the prediction accuracy of AiPPA plays an important role in the design of affinity variants, its performance needs to be improved in the future. However, to improve their performance, deep learning models typically require training with a large amount of high-quality data. Since experimental measurement of the protein-protein binding affinities is very expensive, current experimental affinity data are not sufficient yet to train deep learning models. For example, the PDBbind database, the largest collection of experimental affinity data, contains only about 2,800 samples^38^. Thus, small training datasets remain a challenge for the development of accurate prediction models of the protein-protein binding affinities. Fortunately, with the advances in deep learning, new artificial intelligence strategies such as transfer learning and pre-training have shown promise in improving prediction accuracy and generalization for the models trained on small datasets, providing a potential avenue for future optimization of AiPPA. In recent years, various protein design methods have been used to design or optimize antibodies, including nanobodies^32,54,55^. In contrast to general protein design, antibody design often focuses on the design of the CDR loop sequences and does not require extensive design of their frameworks. In practice, to minimize development risk, a therapeutic antibody usually prefers to have a humanized framework and, as far as possible, its sequence is not altered during the antibody design process. Therefore, the method developed in this study is particularly suited to such an antibody design scenario. In nature, our method mainly designed the sequences of the CDR loops of the template nanobody. Unlike many point mutation optimization methods where the lengths of the CDR loops were fixed^32,55^, our design method allowed changes not only in the amino acid types of the CDR loops but also in their lengths during the MC sampling process of the affinity variants. Therefore, our method can sample a much larger CDR sequence space compared to the point mutation methods. Of course, our method can also be applied to the design of a nanobody framework by defining it as the target design region. Also, given a specific template protein, the method can be easily extended to design other proteins, including the traditional full-length antibodies. Therefore, our computational design method presented in this work is a novel approach to antibody design.

In summary, while nanobodies represent a promising class of protein therapeutics, the experimental screening of high-affinity nanobodies against specific antigens remains time-consuming and resource-intensive. To address this challenge, we developed a novel computational framework that integrates GNNs with MC sampling to enable efficient design of affinity variants of a given antigens. By applying this physics-informed deep learning approach, we successfully designed two nanobodies targeting TL1A, a key therapeutic antigen in autoimmune diseases. This study not only provides a novel development strategy for protein therapeutics but also offers novel antibody molecules with therapeutical potential for the treatment of autoimmune diseases.

## Methods

### Construction of protein graph

Each protein in a protein-protein interaction pair is represented as an undirected graph *G*(*V*, *E*, *A*), where *V* denotes the set of nodes, each representing a residue in the protein; *A* is the adjacency matrix of the graph with dimensions *N*_*v*_ × *N*_*v*_; and *E* = {*e*_*vw*_ | *A*_*vw*_ = 1} is the set of edges in the graph. An edge only exists between node *v* and node *w* if and only if their Cα-Cα distance is less than or equal to the threshold *θ*:

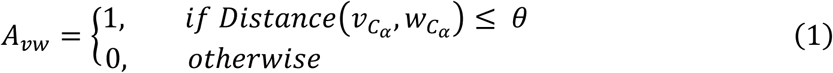

We used the residue type (*AAtype*_*v*_), residue-level solvent-accessible surface area (*SASA*_*v*_ ), and Cα coordinates 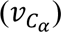 as the initial features of each node. The 20 amino acid types were encoded as integers ranging from 0 to 19 and transformed into 10-dimensional vectors. The *SASA*_*v*_ values were directly calculated using Biopython (https://biopython.org) based on the protein structure. Instead of directly using the Cα coordinates of the residues, we first translated the protein such that its centroid was positioned at the origin. The transformed Cα coordinates were then used as the positional features 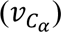. Finally, these three features were concatenated to form the initial node features:

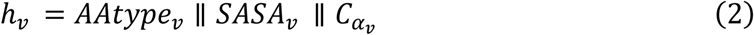

where || represents concatenation operator, resulting in an initial feature dimension of 14 for each node.

We used two attributes as the initial edge features: the distance between nodes (Distance) and the code describing whether the corresponding residues belong to the same chain (EdgeType). The distance between two residues was measured by the Euclidean distance between their Cα atoms. EdgeType was represented as 0 or 1, indicating whether the residues were in the same chain. EdgeType was further transformed into a 7-dimensional vector. Consequently, the initial edge dimension was 8:

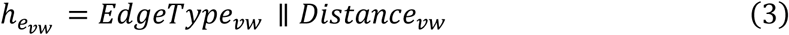

### Architecture of the AiPPA model

During the node and edge updates at layer *t*, we first aggregate the edge features as “message”:

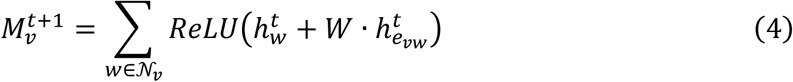

where 𝒩_*v*_ represents all the neighboring nodes of node *v* and *W* is a learnable weight matrix. In the first layer, the hidden states of the nodes and edges are their initial features. The message is then passed to the node *v*, and update its hidden state using the GINE model:

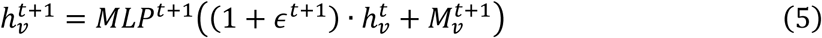

where *ϵ* is a parameter that is learnable. Next, batch normalisation (BN) is performed and then nonlinear activation is applied to update the hidden states:

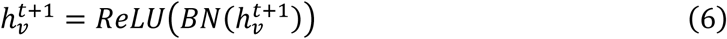

After updating the nodes, pass their hidden states to the adjacent edges to update the hidden states of the edges:

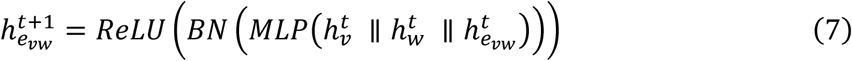

After stacking two EGINE layers ( *t* = 1, 2 ), “global add” operation is performed to aggregate the features of all the edges in the last layer, producing the final representation of the protein graph:

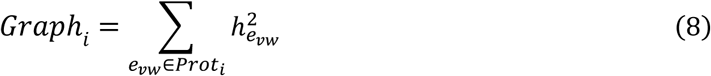

where *i* = *A or B*, and represents each of the two interacting proteins. Finally, the representations of the two interacting protein graphs are combined and fed into a three-layer MLP to predict the binding free energy of the two proteins (A and B):

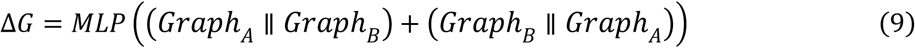

### Monte Carlo sampling of low-BFE variants

To generate CDR sequences of a given variant by the MC sampling, three operations with different probabilities were used to explore the sequence space: point mutation, insertion, and deletion. Point mutation (length-preserving mutations) replaces an amino acid at a given position with different type based on a specific probability; and insertion and deletion (length-altering mutations) modify the CDR lengths by randomly adding or removing an amino acid.

The probabilities of these operations were similar to the study by Yeh et al.^56^: point mutations were assigned a probability of 70%, and insertions and deletions together accounted for 30%.

Based on the CDR length distributions of the natural nanobodies, the CDR length ranges for sampling are: CDR1 - 7 amino acids, CDR2 - 5 or 6 amino acids, and CDR3 – randomly varying from 5 to 20 amino acids. The probabilities for insertion and deletion (length transition probabilities) depended on the length of a given CDR: deletion (eq. 10) and insertion (eq. 11) probabilities were determined by the proportion of sequences with the same CDR length observed in the natural nanobodies:

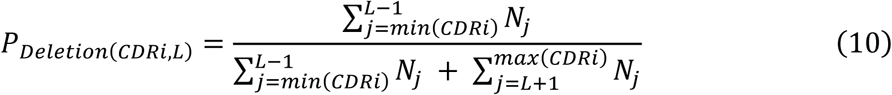

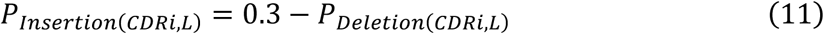

where *i* = 2 *or* 3, L is the given length of CDR2 or CDR3, and *N_j_* is the number of the corresponding CDRs with length *j* in the INDI database. For insertion and point mutation, we used the amino acid composition at each CDR position of the natural nanobodies as the sampling probability. In addition, for the hypervariable region of CDR3 (100A–100L), we used the amino acid frequencies given in the McMahon library^57^ as the sampling probability.

After calculating the binding free energy (Δ*G*) between a given variant and the protein target using AiPPA, its acceptance or rejection was determined by the Metropolis sampling criterion:

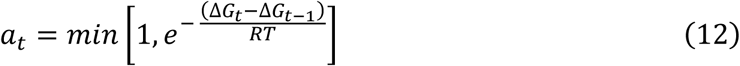

where *a*_*t*_ is the acceptance probability of step *t*, Δ*G*_*t*_ is the binding free energy predicted by AiPPA at the step *t*, and Δ*G*_*t*−1_ is the binding free energy at the previous step *t*-1; *R* is the ideal gas constant of 8.314 J mol⁻¹ K⁻¹; and *T* is temperature. In this study, we set *T* to the room temperature times 0.2 (i.e., 59.6 K) in order to slightly increase the acceptance probability of the variants with higher binding free energies.

### Local refinement of binding interface

The binding interface residues of a given nanobody were defined as those with at least one heavy atom within 4.5 Å of TL1A. To refine the interface residues, all non-interface residues in the given nanobody were fixed. Then, ProteinMPNN (version: v_48_020.pt) was run using two sampling temperatures (0.1 and 0.3) to generate 100 interface sequences for each temperature, with the following parameters:

> *python protein_mpnn_run.py \*
>
> *--jsonl_path ../parsed_pdbs_bb.jsonl \*
>
> *--chain_id_jsonl ../assigned_chains.jsonl \*
>
> *--fixed_positions_jsonl ../masked_pos.jsonl \*
>
> *--out_folder $output_dir \*
>
> *--num_seq_per_target 100 \*
>
> *--sampling_temp “0.1 0.3” \*
>
> *--seed 37 \*
>
> *--batch_size 1*

where the file *parsed_pdbs_bb.jsonl* contains PDB information of the given structure, *assigned_chains.jsonl* specifies chain details, and *masked_pos.jsonl* defines the fixed positions, allowing for targeted site design. These three files were generated by default.

After the local refinement using ProteinMPNN, we evaluated the binding affinities of the refined sequences also using a method described by Yi et al.^53^, which strongly correlates with the presence of six interface residues: tyrosine (Y), glycine (G), serine (S), arginine (R), valine (V), and isoleucine (I). Given that ProteinMPNN generates sequences with fixed backbones, we assumed the refined sequences would maintain the original complex structures, preserving the positions of interface residues. Based on the initial complex structures, we analyzed the interface residues of the refined sequences, calculated the affinities, and then selected the top 10 sequences with the highest predicted affinities for further interface quality analysis.

### Protein expression

The nanobody gene was inserted into the pET26-b (+) expression vector, incorporating a pelB signal peptide at the N-terminus, a TEV protease cleavage site, and a 6×His tag at the C-terminus. The expression vector was transformed into *E. coli* Rosetta (DE3), and the cells were cultured in LB medium at 37°C until the OD600 reached 0.5–0.6. Gene expression was induced with 0.1 mM IPTG and allowed to proceed for 20 hours at 25 °C. Harvested cells were resuspended in SET buffer (30 mM Tris-HCl, 1 mM EDTA, 20% sucrose, pH 8.0) and incubated on ice for 20 minutes. After centrifugation at 8000 × g for 20 minutes at 4 °C, the cell pellet was resuspended in pre-cooled 5 mM MgSO4 and incubated on ice for a further 20 minutes.

The periplasmic fraction was separated from the cell debris by centrifugation at 11,000 rpm for 20 minutes at 4 °C, and the supernatant was collected.

### Protein purification

One mL of Ni-NTA Beads 6FF resin was packed into a pre-assembled chromatography column (BIO-RAD) and equilibrated with 10 mL of buffer A1 (500 mM NaCl, 10 mM Na₂HPO₄, 10 mM NaH₂PO₄, 20 mM imidazole, pH 7.4). The supernatant obtained from the protein expression step was then loaded onto the column. Afterward, the column was washed with 10 mL of buffer A1 to remove non-specifically bound impurities. Subsequently, the bound proteins were eluted with 5 mL of elution buffer (500 mM NaCl, 10 mM Na₂HPO₄, 10 mM NaH₂PO₄, 300 mM imidazole, pH 7.4), and the eluate was collected for the downstream processing.

The eluted protein fraction was dialysed overnight at 4°C using Yuanye cellulose dialysis bags (MWCO 8000-10000, catalogue number SP131264) against 1× PBS as the dialysis buffer. Following dialysis, the protein was treated with TEV protease (Smart-Life-Sciences) at 25°C for 2 hours to cleave the His tag. The tag-cleaved protein was then subjected to reverse affinity chromatography on a pre-equilibrated Ni-NTA column in PBS buffer to remove any residual His-tagged contaminants. Finally, the purified tag-free protein was concentrated using a 10 kDa MWCO centrifugal filter device (Amicon), aliquoted as required, and rapidly frozen in liquid nitrogen. The resulting protein was stored at -80 °C for long-term storage.

### Bio-layer interferometry measurements

Bio-layer interferometry (BLI) measurements were performed on the Sartorius Octet R8 protein analysis system using Octet® SA biosensors purchased from Sartorius. BLI experiments were performed at 25°C using PBS (pH 7.4) supplemented with 0.02% (w/v) Tween-20 as the detection buffer. Samples were prepared in a concentration gradient, and a control group was included to account for non-specific binding.

A BLI measurement began with sensor equilibration in detection buffer for 90 seconds. Biotinylated TL1A was then immobilized on the biosensors for 300 seconds until a binding signal of 3.0 nm was achieved. The sensors were then equilibrated in the detection buffer for 180 seconds. The sensors were then exposed to wells containing Br3s95 (5,000, 4,000, 3,250, 2,500, 1,250, 625 and 312.5 nM) and Tr3s90 (6,000, 5,000, 4,000, 2,500, 1,250, 625 and 312.5 nM) nanobodies for 160 seconds to allow binding. Finally, the sensors were immersed in detection buffer for 100 seconds to monitor dissociation.

The BLI data of the protein-protein interaction kinetics were analyzed using the Octet® Analysis Studio Software (ver. 13.0.1.35). Non-specific binding was corrected by subtracting signals from the control group blank wells and the binding curves were fitted to the 2:1 heterogeneous binding model because of multiple affinity binding sites on the target protein.

### Data availability

All datasets used for the model training and evaluation are publicly available, with the processing details described in Methods and Supplementary Methods.

### Code availability

The source codes for AiPPA are publicly available via GitHub at https://github.com/Fudan-HQLab/AiPPA.

## Acknowledgements

We think Dr. Yong Zhao (U-mab Biopharma, Inc.) for his valuable assistance during the initial phase of this project. This work was supported by the National Key Research and Development Program of China (2021YFA0910604), and the National Natural Science Foundation of China (31971377, 31671386).

## Author contributions

Q.H. supervised the project. L.W., X.H. and Q.H. conceived the project; L.W. developed the computer programs and performed the computational study; L.W., G.G. and X.Q. collected the data and analyzed the computer codes. X.H. and G.G. performed the experimental validation and analysis; L.W. and Q.H. drafted and revised the manuscript. All authors contributed to the drafting and revision of the paper.

## Additional information

**Supplementary information** is available for this paper at …

**Correspondence and requests for materials** should be addressed to Q.H.

