## Supplementary Information for "Computational nanobody design using graph neural networks and Metropolis Monte Carlo sampling"

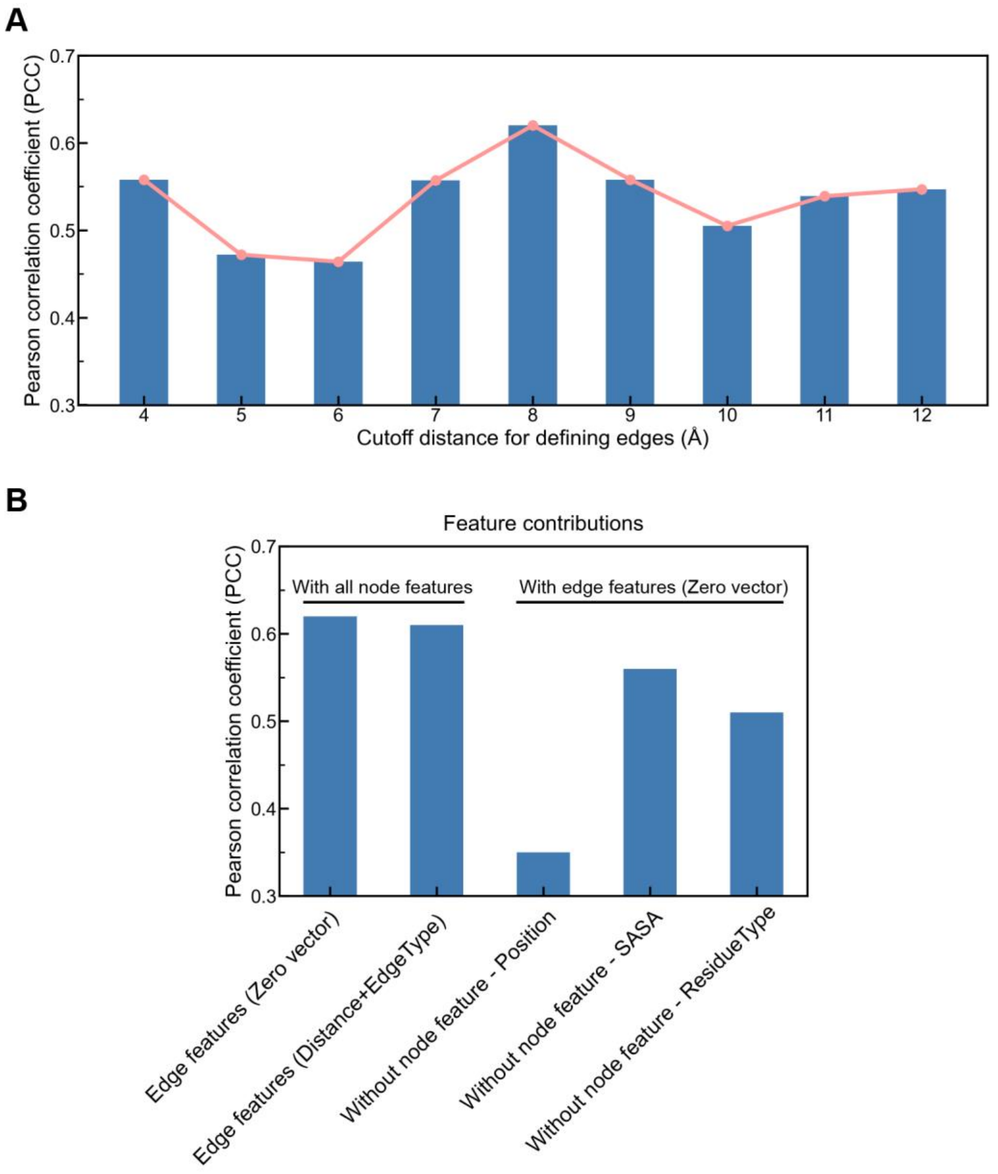


**Figure S1. Impact of cutoff distance and initial features on the performance of the AiPPA model.** (A) Performance of AiPPA with different cutoff distances for edge definition. Optimal performance was achieved when edges were defined between nodes with a Cα–Cα distance ≤ 8 Å. (B) Performance of AiPPA with different feature combinations. Using zero vectors as initial edge features outperforms the initial non-zero features (Distance + EdgeType). With zero vectors as initial edge features, the contributions of three node features to the performance were different.

### Supplementary Methods

(1) *Datasets for training and validation*

The PDBbind dataset (v2020) provided raw structural files identified by PDB codes. Crystal structures often contain multiple polypeptides or proteins, and X-ray diffraction experiments may yield multiple copies of interacting proteins. For binding affinity analysis, the biologically relevant conformation—known as the biological assembly—is typically used (Supplementary Fig. 2). For example, PDB 7RBY includes two biological assemblies, each comprising the RBD domain of the SARS-CoV-2 Spike protein and a nanobody (Nb112). For PDBbind structures with multiple biological assemblies, one assembly was randomly selected for analysis.

After excluding entries present in the test set (described below), we applied a screening and processing workflow to the remaining PDBbind entries, resulting in 1,858 entries for model training and validation. The processing steps included:

1. **Selection of entries**: Entries with specific $K_\mathrm{i}$ or $K_\mathrm{d}$ values were selected.
2. **Backbone completion**: Missing heavy atoms in the backbones were added.
3. **Residue standardization**: Non-standard residues were converted to their standard equivalents.
4. **Solvent and heteroatom removal**: Solvent molecules, small molecules, and other heteroatoms were deleted.
5. **Chain selection**: Chains with at least 30 residues were retained.

(2) *Dataset for evaluation*

We applied the same screening and processing workflow used for PDBbind entries to the Kastritis benchmark dataset, and generated a test set of 138 samples for model evaluation.

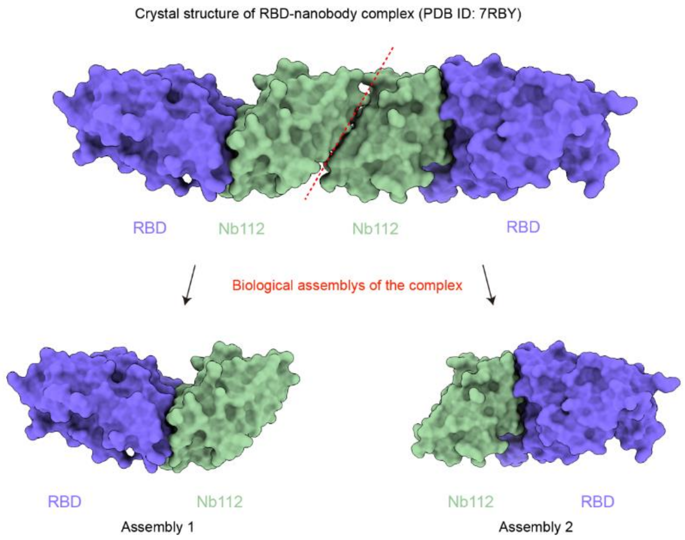


**Figure S2. Crystal structure with multiple biological assemblies.** The complex structure of nanobody Nb112 bound **to** SARS-CoV-2 Spike RBD with two biological assemblies (PDB ID: 7RBY)**.** One assembly was selected randomly for the training set inclusion.

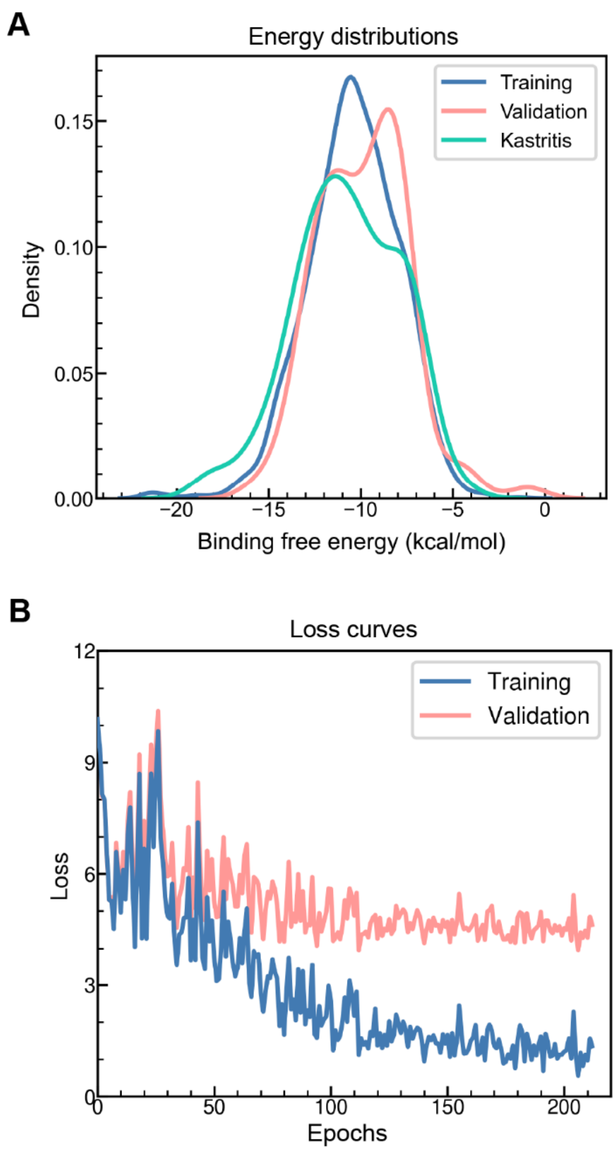


**Figure S3. Training of AiPPA. a**, Distributions of binding free energies across the training, validation, and test sets. Binding free energies span a wide range with similar distributions in all three sets. **b**, Loss (mean squared error, MSE) curves for the training and validation sets. AiPPA loss values consistently decrease and converge across both sets.

**Table S1. The performance of the AiPPA model.**

| Models | Kastritis benchmark | | Rigid complexes | | Flexible complexes | |
|---|---|---|---|---|---|---|
| | PCC | RMSE ($kcal·mol^{-1}$) | PCC | RMSE ($kcal·mol^{-1}$) | PCC | RMSE ($kcal·mol^{-1}$) |
| PPA_Pred2 | 0.56 | 2.56 | 0.57 | 2.58 | 0.55 | 2.54 |
| PRODIGY | 0.49 | 2.63 | **0.61** | **2.36** | 0.43 | 2.86 |
| PPI-Affinity | 0.54 | **2.39** | 0.53 | 2.53 | 0.55 | **2.26** |
| AiPPA (***b***) | 0.61 | 2.49 | 0.57 | 2.68 | **0.67** | 2.30 |
| AiPPA (***ub***) | **0.62** | 2.50 | 0.58 | 2.69 | **0.67** | 2.32 |

Note:

(1) ***b*** (bound state): This refers to calculations where the monomer conformations were taken from the complex in their bound state.

(2) ***ub*** (unbound state): This refers to calculations where the monomer conformations are taken from their free, unbound state.

(3) The **bold** formatting numbers are used to highlight the best results.

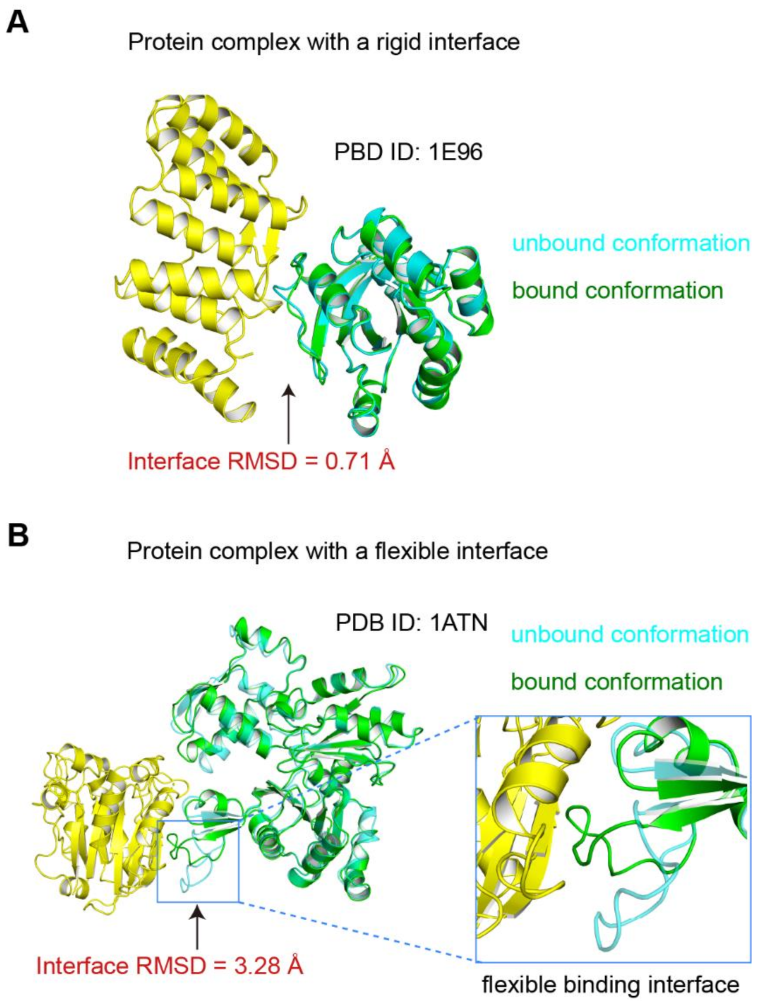


**Figure S4. Protein-protein interaction interfaces of rigid and flexible complexes.** (A) In rigid complexes, interfacial amino acids undergo minimal conformational changes upon binding (Interface RMSD ≤ 1 Å). The unbound protein conformation (cyan) closely resembles the bound state (green). (B) In flexible complexes, interfacial amino acids exhibit significant conformational changes (Interface RMSD > 1 Å). The unbound protein (cyan) often requires specific conformational transitions (green) to form a stable complex. As the AiPPA model predicts binding free energy based on monomer conformations, the use of the unbound state as the input structure is particularly critical.

**Table S2. Insertion codes at the CDR positions.**

| CDR loop | Length | Insertion code | Left | Right |
|---|---|---|---|---|
| CDR1 | 7 | None | 26-27-28-29-30-31-32 | |
| CDR2 | 5 | None | 52 | 53-54-55-56 |
| | 6 | 52A | 52 | 53-54-55-56 |
| CDR3 | 5 | None | 95-96-97 | 101-102 |
| | 6 | None | 95-96-97-98 | 101-102 |
| | 7 | None | 95-96-97-98-99 | 101-102 |
| | 8 | None | 95-96-97-98-99-100 | 101-102 |
| | 9 | 100A | 95-96-97-98-99-100 | 101-102 |
| | … | … | 95-96-97-98-99-100 | 101-102 |
| | 20 | 100A~100L | 95-96-97-98-99-100 | 101-102 |

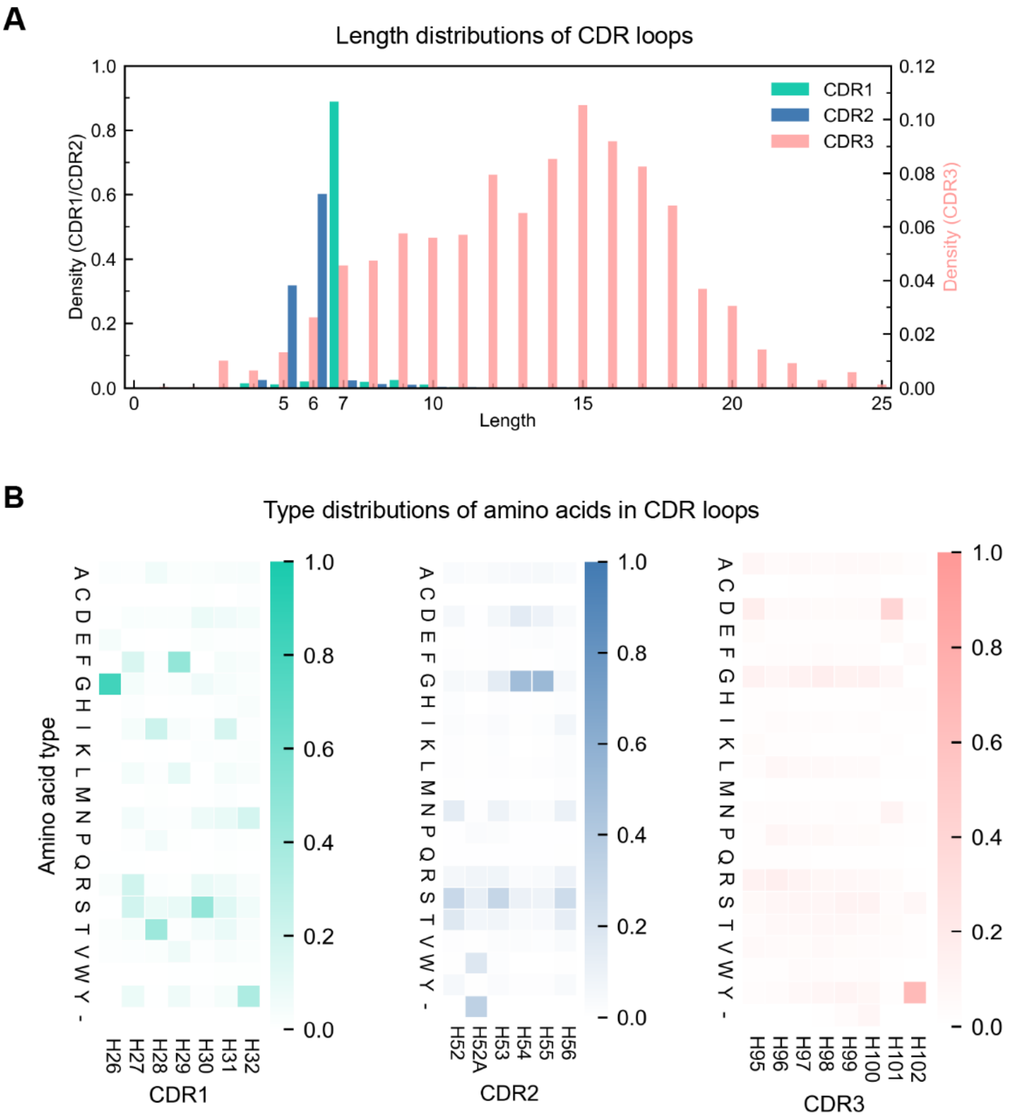


**Figure S5. Sequence characteristics of natural nanobody CDR loops.** (A) Length distributions of natural nanobody CDR loops in the INDI database. CDR1 and CDR2 predominantly span 7 and 5–6 residues, respectively. In contrast, CDR3 exhibits a broader length distribution, with ~95% of sequences ranging from 5 to 20 amino acids. (B) Heatmap of amino acid type distributions at key positions in natural nanobody CDR loops. Certain positions show notable conservation and a preference for specific residues, such as G at H26 in CDR1 and Y at H102 in CDR3. Charged amino acids in CDRs are predominantly D and R.

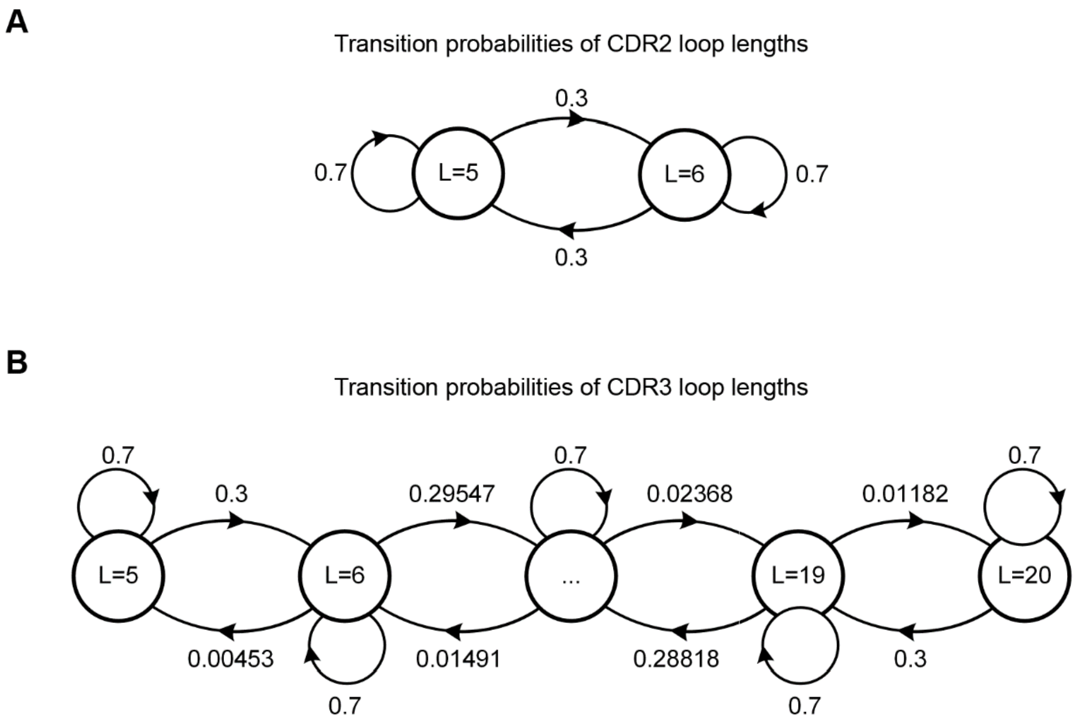


**Figure S6. Transition probabilities of CDR lengths during variant sampling.** (A) CDR2 length transition probability. CDR2 transitions between 5 and 6 amino acids with equal probability, while retaining its current length with a probability of 0.7. (B) CDR3 length transition probabilities. CDR3 length varies continuously between 5 and 20 amino acids, with distinct probabilities for each transition. CDR3 length also maintains its current length with a probability of 0.7.

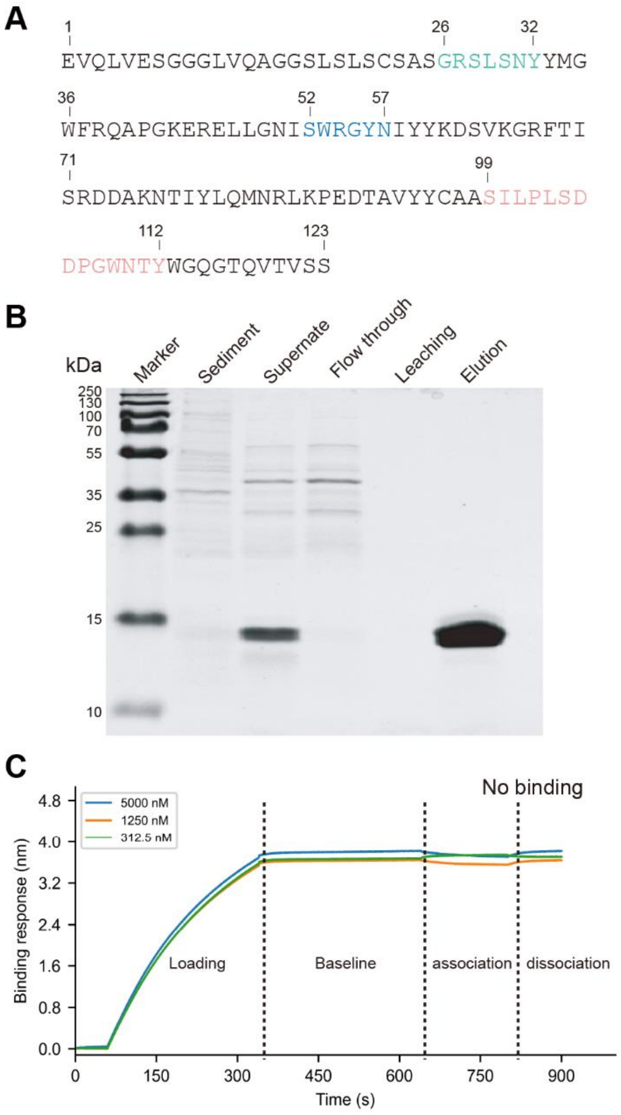


**Figure S7. The template nanobody VHH1.** (A) Amino acid sequence of nanobody VHH1. CDR1, CDR2, and CDR3 are highlighted in cyan, blue, and pink, respectively. (B) SDS-PAGE analysis of expressed nanobody VHH1. (C) BLI measurements of binding kinetics between VHH1 and TL1A.

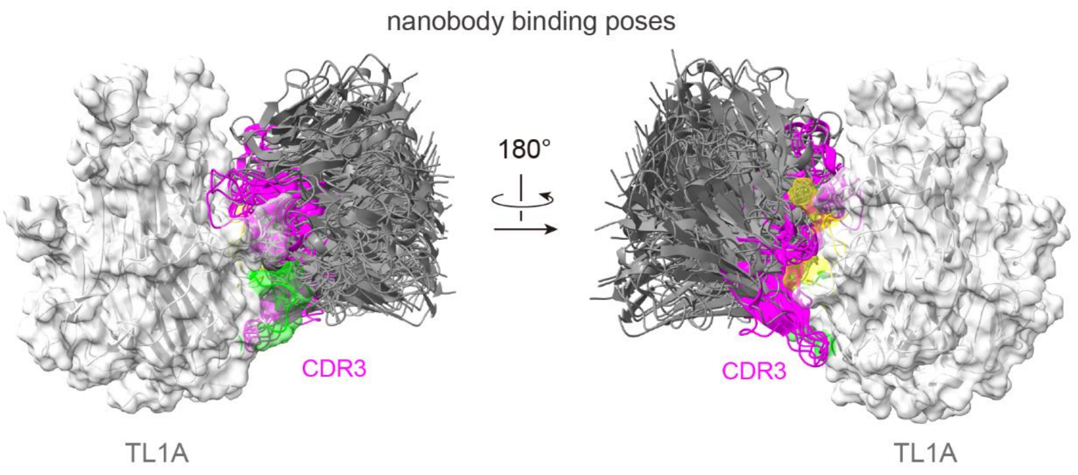


**Figure S8. Examples of HDOCK best-docking poses of designed nanobodies to TL1A.** The desired binding epitope of TL1A is highlighted in lime and yellow.

### Table S3. Site-specific filtering results of the designed nanobodies.

| Design | Solubility (≥ 0.45) | pLDDT CDR1 | pLDDT CDR2 (≥ 80) | pLDDT CDR3 | InterfaceAnalyzer dG_separated (< -40) | InterfaceAnalyzer dG/dSASA×100 (< -1.5) | InterfaceAnalyzer packstat (> 0.65) |
|---|---|---|---|---|---|---|---|
| Br3s56 | 0.50 | 91.75 | 98.02 | 89.36 | -64.03 | -2.71 | 0.66 |
| Br3s95 | 0.56 | 92.61 | 98.09 | 90.88 | -64.01 | -2.71 | 0.70 |
| Br4s38 | 0.50 | 93.15 | 98.10 | 90.67 | -59.05 | -2.46 | 0.67 |
| Tr2s81 | 0.60 | 90.48 | 96.82 | 85.15 | -63.33 | -2.80 | 0.60 |
| Tr3s90 | 0.54 | 89.50 | 96.15 | 85.56 | -61.47 | -2.44 | 0.67 |
| Tr4s18 | 0.45 | 91.65 | 96.77 | 86.69 | -59.09 | -2.85 | 0.67 |

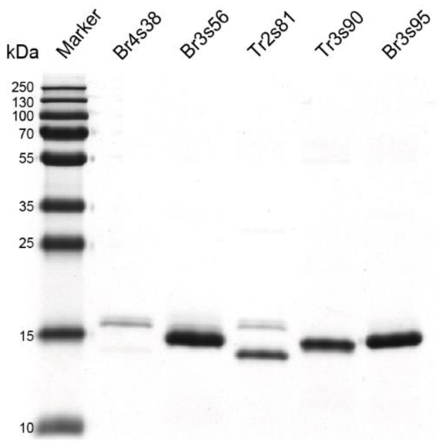


**Figure S9. SDS-PAGE analysis of expressed nanobodies**. Except for Tr4s18, the other five designed nanobodies were successfully expressed in the soluble form.

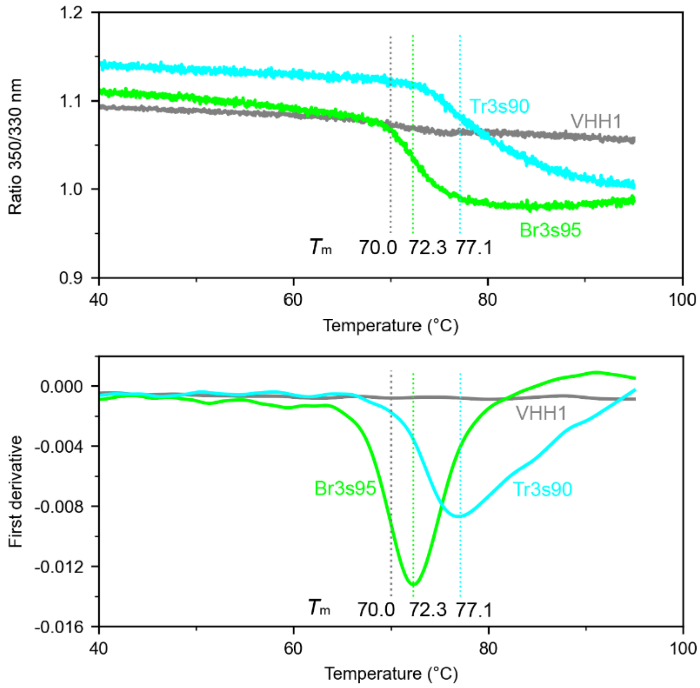


**Figure S10. Thermal stability analysis by Nano-differential scanning fluorimetry (nanoDSF).** Denaturation curves of three nanobodies. The first derivatives of the curves indicate the melting temperatures ($T_m$) of the nanobodies as: VHH1: 70.0°C; Br3s95: 72.3°C; Tr3s90: 77.1°C.
